# FRONTAL EYE FIELD INACTIVATION DIMINISHES SUPERIOR COLLICULUS ACTIVITY, BUT DELAYED SACCADIC ACCUMULATION GOVERNS REACTION TIME INCREASES

**DOI:** 10.1101/187492

**Authors:** Tyler R. Peel, Suryadeep Dash, Stephen G. Lomber, Brian D. Corneil

## Abstract

Stochastic accumulator models provide a comprehensive framework for how neural activity could produce behavior. Neural activity within the frontal eye fields (FEF) and intermediate layers of the superior colliculus (iSC) support such models for saccade initiation, by relating variations in saccade reaction time (SRT) to variations in parameters such as baseline, rate of accumulation of activity, or threshold. Here, by recording iSC activity during reversible cryogenic inactivation of the FEF in non-human primates, we causally test which parameter(s) best explains concomitant increases in SRT. While FEF inactivation decreased all aspects of ipsilesional iSC activity, decreases in accumulation rate and threshold poorly predicted accompanying increases in SRT. Instead, SRT increases best correlated with delays in the onset of saccade-related accumulation. We conclude that FEF signals govern the onset of saccade-related accumulation within the iSC, and that the onset of accumulation is a relevant parameter for stochastic accumulation models of saccade initiation.

**Significance Statement:** The superior colliculus (SC) and frontal eye fields (FEF) are two of the best-studied areas in the primate brain. Surprisingly, little is known about what happens in the SC when the FEF is temporarily inactivated. Here, we show that temporary FEF inactivation decreases all aspects of functionally-related activity in the SC. This combination of techniques also allowed us to relate changes in SC activity to concomitant increases in saccadic reaction time (SRT). Although stochastic accumulator models relate SRT increases to reduced rates of accumulation or increases in threshold, such changes were not observed in the SC. Instead, FEF inactivation delayed the onset of saccade-related accumulation, emphasizing the importance of this parameter in biologically-plausible models of saccade initiation.

## Introduction

How does the brain commit to a voluntary action? The oculomotor system that moves our line of sight provides a model system for study of this question. The primate frontal eye fields (FEF) and intermediate layers of the superior colliculus (iSC) are two of the most studied oculomotor structures, and within each, saccade-related activity peaks around the time of saccade initiation (see Gandhi and Katnani, 2011; Schall, 2015 for review). Such patterns conform well to stochastic accumulator models relating neural activity to saccade initiation (Hanes and Schall, 1996; Dorris et al., 1997; Paré and Hanes, 2003; Ratcliff et al., 2003; Ding and Gold, 2012). Despite this work, simple yet fundamental questions about the oculomotor system remain unresolved. For example, what happens to iSC activity when FEF is suddenly compromised, and how do changes in iSC activity relate to concomitant increases in saccadic reaction times (SRTs) (Peel et al., 2014)? Answering these questions would not only advance basic understanding of communication within the oculomotor system, but can also test the neural implementation of stochastic accumulator models for saccade initiation. However, such answers are surprisingly hard to predict for a variety of reasons.

First, these structures are highly interconnected, by virtue of monosynaptic corticotectal projections (Leichnetz et al., 1981; Komatsu and Suzuki, 1985), and polysynaptic descending (e.g., through the basal ganglia, or other cortical structures), ascending (e.g., via the thalamus or pulvinar; (Sommer, 2003; Berman et al., 2009; Crapse and Sommer, 2009)), and callosal pathways (Pandya and Vignolo, 1971); the FEF and iSC also project directly to the brainstem saccadic burst generator (Raybourn and Keller, 1977; Schnyder et al., 1985; Huerta et al., 1986). Second, the functional content of signals relayed from the FEF to the SC, and indeed between many different oculomotor areas, span the sensorimotor continuum (Segraves and Goldberg, 1987; Everling and Munoz, 2000; Sommer and Wurtz, 2000; Wurtz et al., 2001; Helminski and Segraves, 2003). Third, since stochastic accumulator models predict that saccades occur when activity increases above a fixed threshold, then increases in SRT should relate to decreases in the rate at which activity accumulated and/or the baseline level of activity. However, oculomotor thresholds may not be fixed (Jantz et al., 2013), can paradoxically decrease for longer SRTs (Heitz and Schall, 2012), and other parameters such as the onset of accumulation (Pouget et al., 2011), the speed of perceptual evaluation (Shankar et al., 2011), or the time period of integration (Heitz and Schall, 2012) may also impact when the oculomotor system commits to a saccade. Thus, recording iSC activity during inactivation of the FEF can not only address the contribution of the FEF to iSC activity, but can also test whether the observed profiles of iSC activity accompanying increased SRTs match those predicted by contemporary stochastic accumulator models of saccade initiation.

## Materials and Methods

### Subjects and surgical procedures

Four male monkeys (*Macaca mulatta*, monkeys M, G, D, and O weighing 8.7, 11.1, 9.8, and 8.6 kg respectively) were used in these experiments. All training, surgical, and experimental procedures conformed to the policies of the Canadian Council on Animal Care and National Institutes of Health on the care and use of laboratory animals, and were approved by the Animal Use Subcommittee of the University of Western Ontario Council on Animal Care. We monitored the monkeys’ weights daily and their health was under the close supervision of the university veterinarians.

Each monkey underwent two surgeries to permit cryogenic inactivation of the FEF and extracellular recordings from the iSC, In the first surgery, we implanted either unilateral (right side only; monkeys M and G) or bilateral cryoloops into the arcuate sulcus using surgical procedures previously described (Lomber et al., 1999; Peel et al., 2014). Briefly, we performed a small 2.25 cm^2^ craniotomy above the spur of the arcuate sulcus, and implanted two customized, stainless steel cryoloops (each 5-8 mm in length, and 3 mm in depth) into the arcuate sulcus (Figure 1A), which permitted cooling of tissue adjacent to the superior and inferior arms of the arcuate sulcus. Cryoloop temperatures of 3°C silence post-synaptic activity in tissue up to 1.5 mm away without influencing axonal propagation of action potentials (Lomber et al., 1999). Thermal surface imaging of the exposed tissue during surgery revealed that cryoloop cooling did not spread to adjacent gyri. In this manuscript, we only performed unilateral FEF inactivation, and alternated the side of cooling on separate days in monkeys D and O. In the second surgery, we positioned a recording chamber over a 19 mm diameter craniotomy to permit a surface normal approach to either iSC (Rezvani and Corneil, 2008).

**Figure 1.**
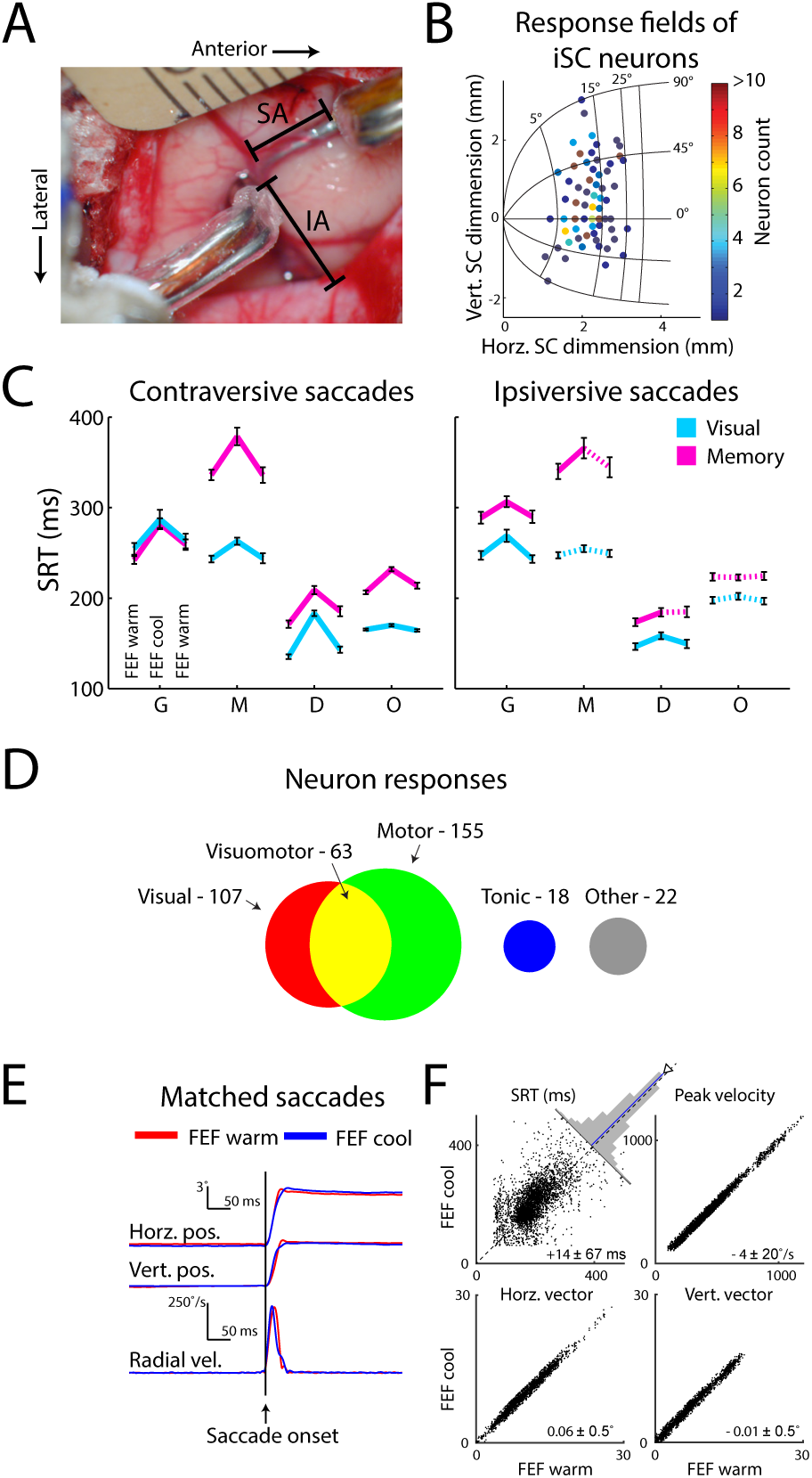
Materials and Methods. (**A**) Cryoloops were inserted into the inferior (IA) and superior (SA) arm of the arcuate sulcus. (**B**) Response field centers for iSC neurons recorded in this study, plotted on the SC map of Hafed and Chen (2016). (**C**) FEF inactivation increased SRTs for contraversive and occasionally ipsiversive saccades. Each line connects mean SRT (+/- SE) across pre-, peri-, and post-cooling sessions for each of the 4 monkeys; solid lines indicate significant differences (p < 0.025, Wilcoxon signed rank test). (**D**) Functional classification of recorded neurons (see Materials and Methods for criteria). (**E, F**) The matched saccade analysis compared saccades of very similar eye position and velocity profiles but different SRTs across the FEF warm or FEF cool conditions (all saccades came from the same session; see Materials and Methods). **E** shows one example of a saccade match; **F** shows characteristics of the 3,762 matched saccade pairs (pooled across both ipsi-and contraversive saccades).

### Experimental procedures

Head-restrained monkeys were placed in front of a rectilinear grid of 500+ red LEDs covering ± 35° of the horizontal and vertical visual field. We conducted experiments in a dark, sound-attenuated room and sampled each monkey’s eye position using a single, chair-mounted eye tracker at 500 Hz (EyeLink II). Behavioural tasks were controlled via customized real-time LabView programs running on a PXI controller (National Instruments) at a rate of 1 kHz.

Extracellular activity was recorded on a MAP data acquisition system (Plexon) via tungsten microelectrodes (impedance 0.5 - 3.0 MΩ at 1 kHz; FHC). Action potential waveforms surpassing a user-defined threshold were amplified, low-cut filtered, sorted, and stored at 40 kHz. All neurons were recorded ~1 mm or more below the surface of the SC, in locations where electrical stimulation (300 Hz, 100 ms, biphasic cathodal-first pulses with each phase 0.3 ms in duration) evoked saccades with currents < 50 μA. In conjunction with most recorded neurons exhibiting delay-or saccade-related activity, recorded neurons were most likely contained with the intermediate, rather than superficial, layers of the SC, but we cannot completely rule out this possibility. We subsequently confirmed the isolation of single-unit neurons offline throughout cooling using both sorted and unsorted action potential waveforms, and when possible ensured that the functional definition of a given neuron was maintained before and after FEF inactivation.

Upon isolating an iSC neuron, we mapped response field for contralateral visually-guided saccades. Across the 239 isolated neurons in our sample, response field centers were located at an eccentricity of 11.6 ± 4.7° (range: 4 to 25°) and an angle relative to the horizontal axis of 12.8 ± 30.5° (range: −90 to 90°) (Figure 1B). After response field mapping, we attempted to collect a dataset consisting of a pre-, peri-, and post-cooling session (60 correct trials each), which required maintaining isolation for ~20 minutes. We largely followed a previously-described procedure for cryogenic FEF inactivation (Peel et al., 2014), although to facilitate neuronal isolation through the entire dataset, we did not implement a 3 minute transition between cooling sessions. Following the completion of the pre-cooling session, chilled methanol was pumped through the lumen of the cryoloops, decreasing the cryoloop temperature. Once the cryoloop temperature was stable at 3°C, we began the peri-cooling session. Upon finishing the peri-cooling session, we turned off the cooling pumps, which allowed the cryoloop temperature to rapidly return to normal. When the cryoloop temperature reached 35°C, we began the post-cooling session. Although saccadic behaviour and iSC activity rapidly recovered after rewarming, the post-cooling sessions may have contained residual effects of cooling. We controlled for this and other time-dependent factors by combining trials from pre-and post-cooling sessions into the FEF warm condition. For ~15% of all datasets, isolation of an iSC neuron was lost after completion of the peri-cooling session. We excluded post-cooling trials from these sessions. Nonetheless, the effects of cooling in these sessions (based on comparing peri-to pre-cooling activity) were similar to the other 85% of datasets where isolation was maintained throughout the post-cool session.

### Behavioural tasks

Monkeys performed visually-or memory-guided saccades after a delayed response period. Following a variable fixation period (750 to 1000 ms) where monkeys maintained fixation within a radius of ±3° of a central cue, a peripheral cue appeared in the periphery. The fixation window was purposely set to be large, as FEF inactivation can shift the average fixation position slightly (as discussed in (Peel et al., 2016), fixation position can shift by ~0.5° on average, although the range of fixation positions adopted before and during FEF inactivation still overlap). In 79% of our datasets, the peripheral cue could appear either in or diametrically opposite to the center of the response field of an isolated SC neuron. In the remaining 21% of datasets, we collected data from intermixed visual-or memory-guided saccades, using peripheral cues placed only in the neuron’s response field. Peripheral cues were either extinguished after 250 ms or remained on for memory-or visually-guided saccades, respectively. To receive a liquid reward, monkeys were required to maintain fixation throughout a delay-period of 1000 ms, and generate a saccade towards a target window (70% of the peripheral cue’s visual angle) when the central cue was extinguished. This large target window was necessary since FEF inactivation increased saccadic error, particularly for memory-guided saccades (Peel et al., 2014).

Consistent with previous reports (Peel et al., 2014, 2016; Kunimatsu et al., 2015), large-volume unilateral FEF inactivation increased contraversive and occasionally ipsiversive saccadic reaction times (Figure 1C), and also decreased the accuracy and peak velocity of contraversive saccades. However, FEF inactivation had only a margin impact on the monkeys’ ability to perform either task, with error rates increasing at most by 14%.

### Neuron Classification

iSC neurons were classified functionally according to a variety of response characteristics on acceptable trials (Figure 1D). To quantify neuronal activity, we convolved spike times on individual trials with a spike density function that mimics an excitatory post-synaptic potential (rise-time of 1 ms, decay-time of 20 ms, kernel window of 100 ms; (Thompson et al., 1996)). We confirmed that all of our results were the same if we convolved neural activity with a 10-ms Gaussian.

For *visual activity*, we defined acceptable trials as those where the monkey maintained fixation of the central cue for the entire delay-period for either visually-or memory-guided saccades, and generated their first saccade towards the target as determined using a velocity criterion of 30°/s. We then checked if neurons exhibited a visually-related response using Poisson analysis described elsewhere (Hanes et al., 1995). Briefly, we compared the actual number of spikes within a time window to the number of spikes predicted by a Poisson distribution based on spiking activity across the entire trial. To calculate the latency of visual response within a trial, we utilized the time of the first burst of spikes greater than chance between 30 to 120 ms after cue onset; the visual latency of a given neuron was then derived by averaging the latency of detected single-trial visual responses across at least 8 trials. In addition, we also ensured that neurons with a visually-related response had mean firing rates in the 50 ms interval after the average visual latency significantly greater than baseline activity integrated in the last 200 ms before cue onset (p < 0.05, Wilcoxon signed rank test; (Basso and Wurtz, 1998; McPeek and Keller, 2002)). Visual activity was defined as the difference between these firing rates. We also calculated the peak magnitude of the visual response minus the baseline activity.

For *delay-period* and *build-up activity*, we applied the same criteria, but also removed any trial with anticipatory saccades (i.e., reaction time less than 60 ms after fixation cue offset; ~12% of trials). Neurons displayed *delay-period activity* if the mean firing rates in the last 100 ms of the delay-period were at least 5 spikes/s above baseline activity (i.e., 200 ms before cue onset; p < 0.05, Wilcoxon signed rank test; (Basso and Wurtz, 1998; McPeek and Keller, 2002)). The magnitude of delay-period activity was then calculated as the difference in firing rates between these intervals. Neurons with *build-up activity* had mean firing rates 100 to 200 ms before saccade onset significantly greater than the preceding 100 ms (p < 0.05, Wilcoxon signed rank test; (Anderson et al., 1998)).

Finally, for *saccadic activity*, we used the same trials as those for the analysis of delay-period and build-up activity, but we additionally removed any trial where the monkey blinked during the first saccade (~11% of all trials in this subset). We subsequently removed any dataset with less than 8 acceptable saccades into the response field of an isolated iSC neuron either before, during, or after FEF inactivation (~9% of all datasets were removed). Neurons exhibited *saccadic activity* if the mean peri-saccadic firing rates (defined as 8 ms before saccade onset to 8 ms prior to its end) were significantly greater than the last 100 ms of the delay-period (p < 0.05, Wilcoxon signed rank test), and if the increase in peri-saccadic activity above baseline activity in the 200 ms before cue onset exceeded 50 spikes/s (Munoz and Wurtz, 1995; McPeek and Keller, 2002). Saccadic activity was defined as the difference of mean peri-saccadic and baseline firing rates. We also calculated the peak magnitude of saccade-related activity minus baseline activity.

### Matched saccade analysis

In order to examine iSC activity associated with saccade initiation and generation across FEF inactivation, it is imperative that the actual saccades being compared are as similar as possible. Otherwise, any differences in saccade-related activity could be due to the generation of a saccade of different metrics, given the spatial coding of saccade metrics in the iSC, or different peak velocity, given potential relationships between the vigor of iSC activity and peak saccade velocity (Waitzman et al., 1991; Stanford et al., 1996; Katnani and Gandhi, 2012). To avoid these confound, for each neuron, we matched each FEF cool trial with one trial from a set of corresponding FEF warm trials containing similar saccade metrics and kinematics (Figure 1E). We specified that any such matched saccades had to have horizontal and vertical displacements within 1.5°, and peak velocities within 50°/s. If a FEF cool trial matched with multiple FEF warm trials, we selected, with replacement, the closest match with the lowest ranked differences in three variables (horizontal and vertical displacements and peak velocities), and occasionally by radial displacement for ties. Importantly, this ranking procedure ensured that most matches usually had differences in horizontal and vertical displacement much less than 1° (mean ± SD of 0.06 ± 0.5° and −0.01 ± 0.5°, respectively), and less than 10°/s in peak velocity (-4 ± 20°/s) (Figure 1F). Using this matching procedure, we were able to match 80% of FEF cool trials with a corresponding FEF warm trial recorded from the same neuron. Unless otherwise noted, we matched saccades for both metrics and kinematics for all analyses on saccade-related activity, and only analyzed saccade-related activity in neurons where we could match at least 5 trials in each of the FEF warm and FEF cool conditions (96% of neurons with saccade-related activity met this criterion). To assess the variability inherent to this procedure with and without FEF inactivation, we also performed the same matched saccade analysis utilizing only FEF warm trials.

### Detection of the onset of saccade-related accumulation

To investigate the neuronal correlates of SRT increases during FEF inactivation, we derived various parameters of a rise to threshold model (i.e., onset of accumulation relative to the go cue, baseline, threshold, accumulation rate) from saccade-related iSC neurons. To detect the onset time of saccadic accumulation on a trial-by-trial basis, we implemented a piecewise two-piece linear regression of pre-saccadic iSC activity, modifying an approach described elsewhere (Cashaback et al., 2013; Goonetilleke et al., 2015). The objective of this analysis was to find the two linear regressions that best fit the convolved iSC activity before saccade onset; the onset of saccade-related accumulation was taken as the spike time closest to the inflection point between these to linear fits. The first linear regression is based on activity from 100 ms before the go cue (offset of fixation cue) to a candidate inflection point; the second linear regression is based on activity from this candidate inflection point to the peak of pre-saccadic activity. Candidate inflection points are tested at each millisecond within this window prior to peak pre-saccadic activity, and the onset of activity coincided with the time of the spike closest to the inflection point that minimized the summed squared error between convolved iSC activity and the two linear regressions. Note that the slope of the first linear regression was not required to be zero, hence it could capture any delay-period or build-up activity the precedes saccade-related accumulation.

Of the 7,524 total trials (3,762 matches) matched from 193 neurons included in this analysis, we discarded a small percentage of trials (16%) that did not reach certain criteria due to the trial-by-trial variability in saccade-related activity. Of these total trials, we removed 7% of trials with no inflection point (i.e. the minimum summed squared error occurred at start or end of testing window), and 6% of trials where accumulation was not maintained until saccade threshold (i.e. activity decreased after the inflection point). Moreover, we also discarded 1 and 2% of total trials for when onset times occurred before or the go-cue or after saccade threshold, respectively. Because we obtained identical results with or without removing a small fraction of trials with onsets before the go-cue, this suggests that anticipation of the go-cue did not influence our results. These criteria left us with a total of 5,420 trials from 2,710 pairs of matched trials (2,083 pairs from ipsilesional iSC, and 627 pairs from contralesional iSC). Importantly, the r^2^ values of 0.63 ± 0.24 (mean ± SD) produced by this two-piece linear regression were substantially larger than those arising from a single linear fit over the same time range (0.34 ± 0.21). Further, the r^2^ values changed by less that 2% during FEF inactivation (0.63 ± 0.25 for FEF warm trials to 0.61 ± 0.25 for FEF cool trials). To ensure that the first linear regression sufficiently captured delay-period or build-up activity prior to the inflection point, we also performed similar analyses using larger window sizes starting before go cue (200, 350, and 500 ms) or convolution functions (e.g., a 10 ms Gaussian). We also tried fitting the activity prior to the inflection point with a quadratic function. All of these alternative analyses produced robust r^2^ values greater than 0.59, and had similarly detected onset of accumulation values (median difference less than 1 ms across all matched trials) compared to the two-piece piecewise linear regression analysis. Although r^2^ values did significantly increase with the quadratic fit (as expected given the use of an additional term), we deemed that fitting the first portion of the data with a quadratic curve was justified only when this increased the r^2^ value by 10% and significantly reduced the residuals (F-test with p < 0.01; see (Nagy and Corneil, 2010)). Using such criteria, a quadratic fit was deemed necessary on only 15% of all matched trials, and changed onset times by only 6 ± 27 ms (mean ± SD) across all matched trials. We also implemented an alternative Poisson-based analysis of burst onset (Hanes et al., 1995) to verify our main results. For this alternative Poisson-based analysis, we only included trials where we could detect a burst of spikes after the go cue and before saccade onset, with the onset time coinciding with the burst of spikes nearest to saccade onset. We obtained similar results using this alternative onset detection method (see Results).

### Determining accumulation rate, baseline and threshold

We also derived the accumulation rate, baseline, and threshold from saccade-related iSC neurons. For the accumulation rate, we simply took the slope of the second linear regression running between the inflection point and the peak pre-saccadic activity. Because this calculation of accumulation rate hinges on this calculation of onset time, we also derived the accumulation rate in another way by finding the slope of the line when activity crossed the 30 and 70% points of peak pre-saccadic activity. This latter calculation of accumulation rate is independent of the two-piece linear regression, but yielded similar results. For baseline activity, we took the average level of activity in the 100 ms window before the offset of the fixation cue. For threshold activity, we calculated the activity 18 to 8 ms prior to saccade onset (Jantz et al., 2013; Johnston et al., 2014), which is based on the minimum amount of time for the iSC to influence the brainstem circuits regulating saccade onset (Miyashita and Hikosaka, 1996).

Finally, we conducted two further analyses to examine how combined changes of the changes in the parameters of SC saccade-related activity related to SRT differences during FEF inactivation. First, we calculated the time to reach threshold as the difference in the threshold and baseline activities divided by the accumulation rate, and directly related this combined measure to SRT differences using similar millisecond units. Second, because these parameters may co-vary, we examined the change in each parameter (onset of accumulation (ΔO), baseline (ΔB), threshold (ΔT), accumulation rate (ΔA)) in isolation by regressing such changes against the timing residuals *(ε)* of a multiple linear regression consisting of all other parameters (Hewitt et al., 2015). This approach removed the variability associated with interactions amongst the other parameters, so that we could examine how well a single parameter directly contributed to SRT differences. For example, to evaluate ΔO independently of ΔB, ΔT, and ΔA, we used Equation 1 to compute the *ε* term, which represents the remaining temporal variability between ΔSRT and model parameters after everything but ΔO is removed:

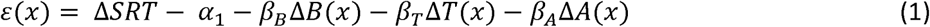

After calculating the timing residuals, we then used a linear regression to fit the final parameter (ΔO) against the *ε(x)* term as shown in Equation 2:

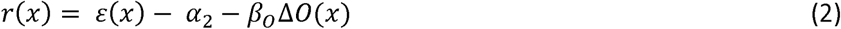

The goodness of fit (r^2^) measure taken from the residuals *r(x)* of this linear regression represents how well changes in onset of accumulation explains SRT differences across matched trials, after removing any variability associated with interactions amongst the other parameters.

### Experimental Design and Statistical Analysis

To quantify the effects of FEF inactivation on visual and delay-period iSC activity, we performed two-sample Wilcoxon rank sum tests to find statistical differences within individual neurons at p < 0.05, whereas we used paired Wilcoxon singed rank tests to uncover statistical differences across the neuronal population at p < 0.05. Because we only analyzed saccadic activity from paired trials having similar saccade metrics and kinematics, we performed paired Wilcoxon singed rank tests on both individual and population of neurons to ensure that differences in saccadic activity reached significance at p < 0.05. We also correlated changes in saccade-related activity during FEF inactivation to concomitant SRT increases. To do this, we performed a regression analysis where we fitted a linear regression to examine how changes in iSC activity predicted SRT differences across the neuron population or all matched trials. We interpreted a significant correlation if the probability associated with the F-statistic (the ratio of the mean regression sum of squares divided by the mean error sum of squares) was less than 0.05.

## Results

We recorded activity from either the ipsi-or contra-lesional iSC before, during, and after large-volume unilateral cryogenic FEF inactivation while monkeys performed delayed visually-or memory-guided saccades to cues placed in or opposite to the center of the neuron&s response field (see Materials and Methods, **and** Figure 1 **for details**). We only cooled the cryoloop in the inferior arm of the arcuate sulcus, which provided an estimated volume of inactivation of 90 mm^3^ in the anterior bank of the arcuate sulcus. Previously, we showed that cooling only the inferior arm cryoloop produced behavioural deficits ~70% of the magnitude produced by cooling both loops together (Peel et al., 2014).

The delayed nature of the behavioral tasks used here, requiring the animals to temporarily withhold a saccade to a persistent or remembered peripheral visual cue until the offset of the central fixation point, provides the opportunity to quantify the effects of FEF inactivation on different aspects of functionally-defined iSC activity. Accordingly, in the first half of the results, we describe the effects of FEF inactivation on iSC activity during visual, delay-period, build-up and saccade-related epochs. In the second half of the results, we focus on how changes in pre-saccade activity in the iSC during FEF inactivation relate to the parameters derived from a stochastic accumulator model. While such models have not customarily been applied to delayed saccade tasks, they provide a framework in which to determine whether changes in baseline, rate of rise, threshold, or onset of accumulation best predicted the associated increases in SRT.

We recorded 239 neurons (178 ipsilesional, 61 contralesional to FEF inactivation) from the caudal iSC, maintaining isolation before and during FEF inactivation, and usually (85%) after re-warming. Of these neurons, 107 (45%) exhibited visual activity, 147 (62%) had delay-period activity, 60 (25%) had build-up activity, 155 (65%) had saccade-related activity, and 22 neurons (9%) did not meet any of our classification criteria. Consistent with previous work (Peel et al., 2014, 2016), unilateral FEF inactivation increased SRT toward contraversive cues, and usually toward ipsiversive cues). Because FEF inactivation also impaired the accuracy and peak velocity of contraversive saccades, we performed a matched saccade analysis to ensure analysis of saccade-related iSC activity during the production of equivalent saccades. Such matching is crucial to avoid confounds related to the generation of a different saccade, or differing degrees of saccade-related drive onto the brainstem burst generator (Yoshida et al., 1999).

### Responses of iSC neurons before FEF inactivation

Our sample of iSC neurons exhibited visual, delay-period, and saccade-related activity, the latencies and magnitudes of which concurred well with previous reports of iSC activity (Munoz and Wurtz, 1995; Basso and Wurtz, 1998; McPeek and Keller, 2002). Before FEF inactivation, neurons with visual activity had response latencies of 56 ± 9 ms (mean ± SD, range: 42 to 99 ms), firing rates of 77 ± 39 spikes/s, and peak magnitudes of 106 ± 54 spikes/s. 80% of these 107 visual neurons also exhibited delay-period activity (71%), build-up (11%), and/or saccade-related activity (59%). Based on these results and our recording approach (see Materials and Methods), we surmise that most visual neurons resided in the intermediate rather superficial layers of the SC. However, the small subset of visual-only neurons (21 total, 14 from the ipsilesional iSC) may have been located in the superficial SC. For neurons exhibiting saccade-related activity, we found firing rates of 160 ± 86 spikes/s and peak magnitudes of 225 ± 108 spikes/s during visually-guided saccades, and corresponding activities of 124 ± 60 and 175 ± 71 spikes/s during memory-guided saccades. 83% of 155 saccade-related neurons also exhibited visual (41%), delay-period (68%), and/or build-up (28%) activity.

We now turn to the effects of FEF inactivation on iSC activity. Where possible, we controlled for time-dependent factors by combining pre-and post-cooling trials into the FEF warm condition, and compared this to trials when the FEF was inactivated (the FEF cool condition). Equivalent results were obtained if we excluded the 15% of our sample where we were not able to record post-cooling data.

### FEF inactivation reduced but did not delay ipsilateral iSC visual responses

We first examine the effect of FEF inactivation on the magnitude and timing of visual responses in the iSC. Figure 2A shows an example neuron recorded from the ipsilesional iSC before, during, and after FEF inactivation. This neuron exhibited delay-period activity since it also remained active at a lower rate when the cue remained on, and also exhibited saccade-related activity (not shown). In the FEF warm condition, visual activity commenced 45 ± 6 ms after cue onset (determined by a Poisson burst analysis; see Materials and Methods), and averaged 71 ± 23 spikes/s for the subsequent 50 ms. FEF inactivation did not alter the visual burst onset latency (45 to 47 ms, p = 0.99, z = −0.01, Wilcoxon rank sum test), but decreased visual activity from 71 to 52 spikes/s (p = 0.06, z = 1.89, Wilcoxon rank sum test).

**Figure 2.**
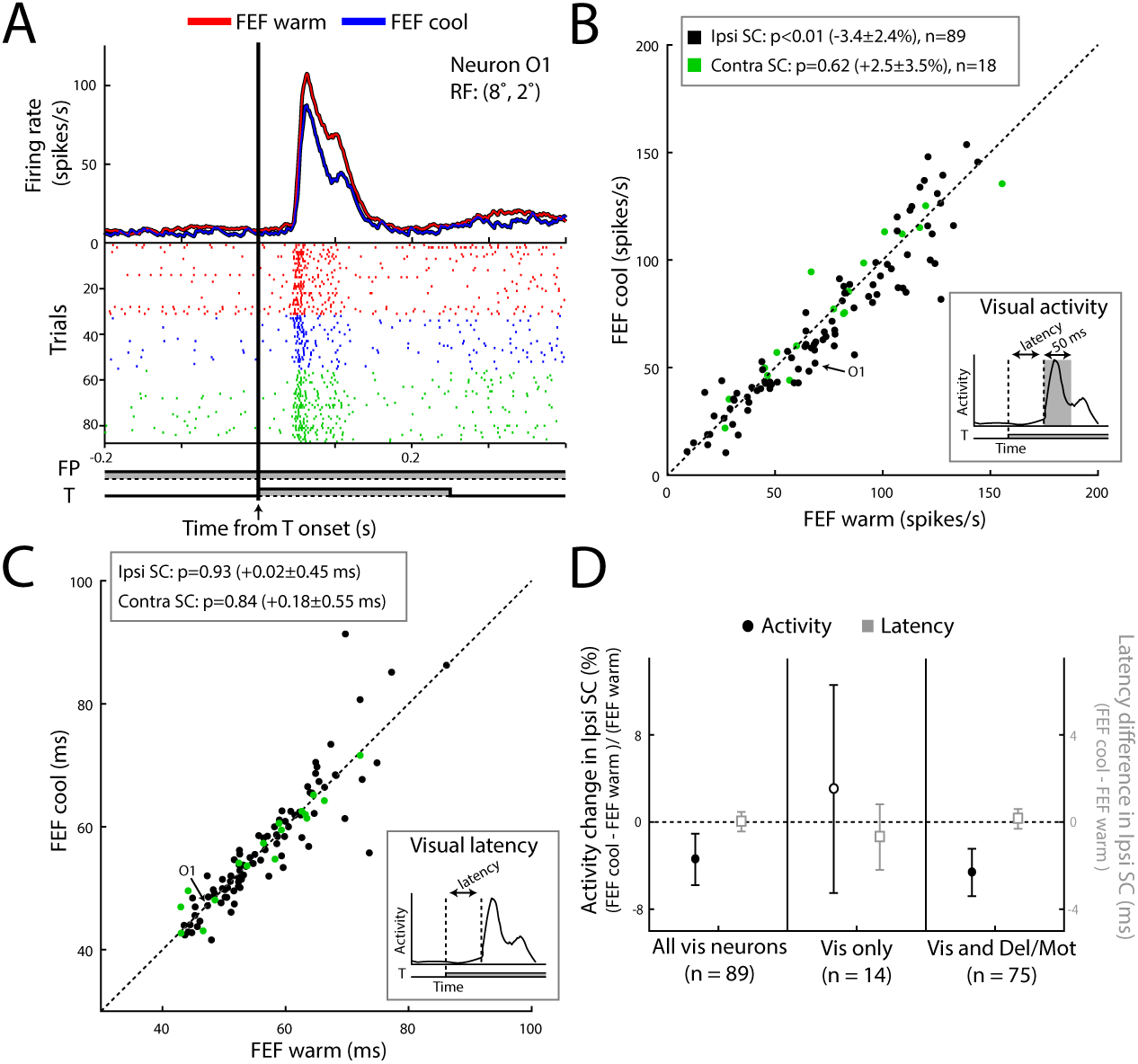
FEF inactivation decreased visual activity of ipsilesional iSC neurons. (**A**) Spike rasters (below) and mean spike density functions (above) showing reduced visual response to peripheral cue onset on ipsilesional iSC neuron O1 with FEF inactivation (FP, fixation point; T, target). (**B, C**) FEF inactivation decreased activity in the 50 ms following the start of the visual response in the ipsi-(black circles) but not contralesional (green circles) iSC (**B**), without altering visual response latency in either iSC (**C**; line represents line of unity; p value shows results of Wilcoxon signed rank test). (**D**) FEF inactivation decreased visual responses of neurons exhibiting other functional responses (left axis, black circles, percent change ± SE), but did not alter visual response latency (right axis, grey squares; difference ± SE). Filled symbols represent significant effects using Wilcoxon signed-rank test (p < 0.05).

Across our sample, FEF inactivation consistently decreased visual activity in the ipsilesional (p < 0.01, z = −2.95, Wilcoxon signed rank test, ~3 ± 2% decrease, mean ± SE, in all 89 neurons, and ~24 ± 3% decrease in the 21 neurons exhibiting significantly decreased visual activity), but not contralesional iSC (3 ± 4% increase; p = 0.62, z = 0.50, Wilcoxon signed rank test, Figure 2). A statistical comparison of changes in the magnitude of the visual response across the ipsi-and contralesional iSC did not reach significance (p = 0.07, z = 1.79, Wilcoxon rank sum test), perhaps because of the small sample size of contralesional iSC neurons. FEF inactivation did not alter visual response latencies across either ipsilesional (increase and SE of less than 1 ms, p = 0.93, z = 0.09, Wilcoxon signed rank test) or contralesional iSC neurons (increase and SE of less than 1 ms, p = 0.84, z = 0.20, Wilcoxon signed rank test; Figure. 2C). When present, FEF inactivation also did not alter activity in the 200 ms preceding cue onset in either the ipsilesional (13 ± 10% increase, p = 0.87, z = 0.16, Wilcoxon signed rank test) or contralesional (2 ± 24% decrease, p = 0.19, z = 1.36, Wilcoxon signed rank test) iSC across our sample, nor in the subset of 21 ipsilesional iSC neurons (7 ± 14% decrease, p = 0.78, z = 0.28, Wilcoxon signed rank test) exhibiting significantly reduced visual activity.

The effect of FEF inactivation on the magnitude of visual responses depended on the functional classification of the recording neuron, being more pronounced in ipsilesional iSC neurons that also displayed delay-and/or saccade-related activity (5 ± 2% decrease, p < 0.01, z = −2.65, Wilcoxon signed rank test, n = 75), compared to the putative superficial SC neurons which only exhibited a visual response (3 ± 10% increase, n = 14; Figure 2D). Although FEF inactivation increased the response magnitude of visual-only neurons, this increase was not significant (p = 0.15, z = −1.48, Wilcoxon signed rank test) given a small sample size and a high degree of variability between neurons. In terms of response latency, and in contrast to the effects of FEF inactivation on response magnitude, we did not observe any influence of FEF inactivation on visual latency of SC neurons when sub-divided into different functional classifications (all differences < 1 ms, p = 0.98 and 0.77, z = −0.03 and 0.28, Wilcoxon signed rank tests on visual only and multiple response neurons, respectively, Figure 2D).

### FEF inactivation decreased delay-period activity in ipsilesional iSC neurons

FEF inactivation reduced delay-period activity in ipsilesional iSC neurons both in the presence (visually-guided saccades) and absence (memory-guided saccades) of peripheral cues. This result is shown for two representative ipsilesional iSC neurons, (Figure 3A&3B). In the memory-guided saccade task, the neuron shown in Figure 3A displayed modest delay-period activity of 13 ± 4 spikes/s in the last 100 ms period before fixation cue offset, which ramped up until saccade onset. FEF inactivation effectively abolished this delay-period activity (decreasing from 13 to 0.2 spikes/s; p < 0.0001, z = 4.90, Wilcoxon rank sum test). The visuomotor neuron shown in Figure 3B was recorded during interleaved visually-and memory-guided saccades, and exhibited far greater delay-period activity when the cue was present (73 ± 6 spikes/s) than absent (12 ± 2 spikes/s). FEF inactivation decreased delay-period activity both when cues were present (73 to 45 spikes/s; p < 0.0001, z = 4.41, Wilcoxon rank sum test) or absent (12 to 5 spikes/s; p < 0.05, z = 2.23, Wilcoxon rank sum test). Across our sample, FEF inactivation robustly reduced delay-period activity in both saccade tasks in the ipsilesional iSC (Figure 3C; visually-guided, cyan, p < 0.0001, z = −4.07, 31% of 77 neurons exhibited significant decreases; memory-guided, magenta, p < 0.0001, z = −4.67, Wilcoxon signed rank test, 40% of 45 neurons exhibited significant decreases). The effects of FEF inactivation were the same on ipsilesional neurons with (squares) or without (circles) build-up activity prior to saccade onset, but proportionally larger during the memory-guided (34 ± 7% decrease in all 45 neurons, or 68 ± 6% decrease in the 18 neurons with significant decreases) versus visually-guided (18 ± 4% decrease in all 77 neurons, or 51 ± 4% decrease in the 24 neurons with significant decreases) saccade task. In contrast, we observed no effect of FEF inactivation on delay-period activity in our sample of contralesional iSC neurons (Figure 3D; p =0.84 and 0.65, z = −0.20 and −0.45, Wilcoxon signed rank tests for memory-guided and visually-guided, respectively). The lack of changes on contralesional iSC neurons during FEF inactivation is also supported by FEF inactivation producing larger decreases of delay-period activity in the ipsi-compared to the contralesional iSC (p values < 0.01 and 0.1, z = 2.76 and 1.37, Wilcoxon rank sum tests for memory-and visually-guided, respectively).

**Figure 3.**
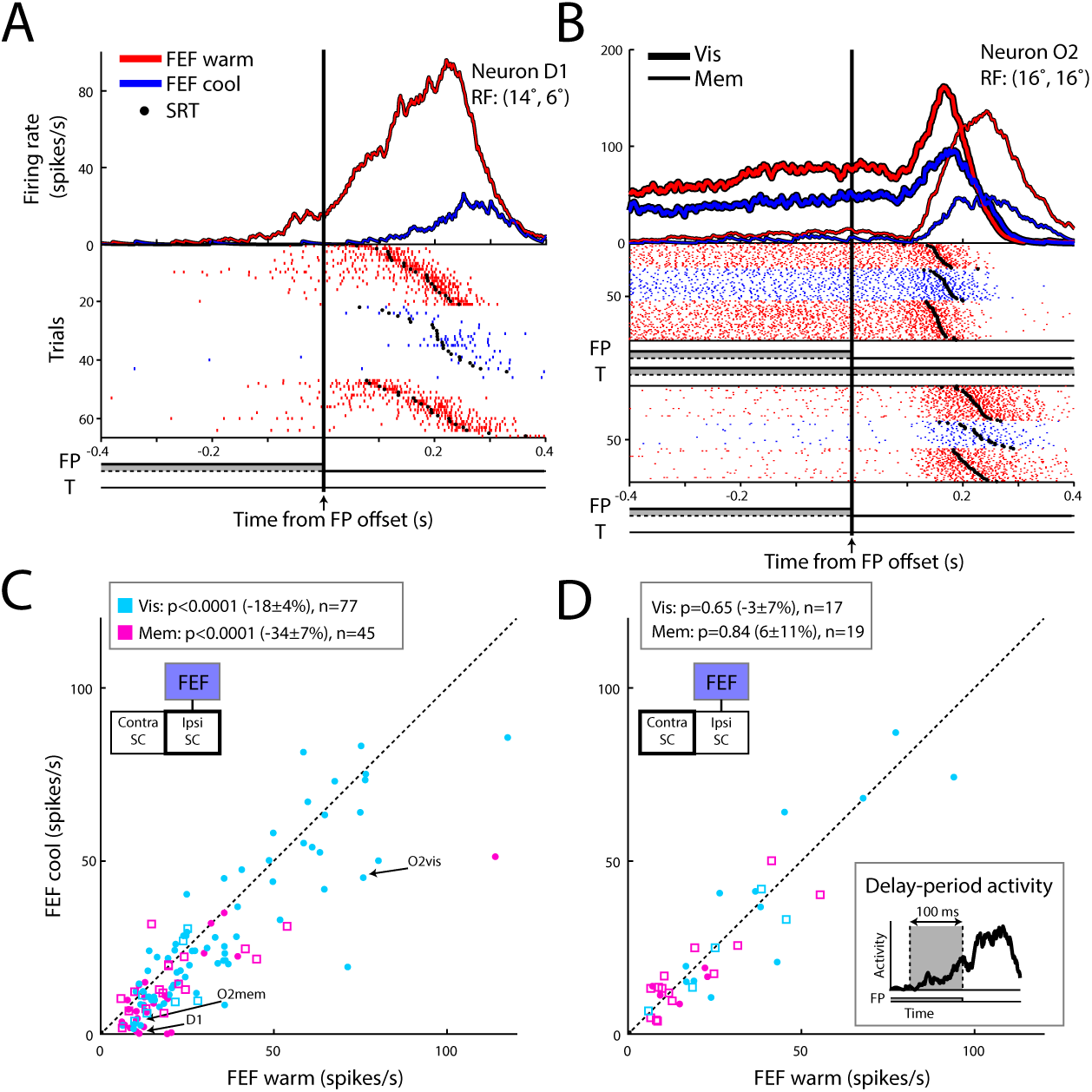
FEF inactivation decreased delay-period activity of ipsilesional iSC neurons. (**A, B**) FEF inactivation nearly abolished modest delay-period activity in ipsilesional iSC neuron D1 during a memory-guided saccade (**A**), and reduced delay-period activity in ipsilesional iSC neuron O2 in both the visually-and memory-guided tasks (**B**). (**C, D**) FEF inactivation consistently decreased delay-period activity in the last 100 ms before peripheral cue offset for both visually-and memory-guided tasks in ipsilesional (**C**) but not contralesional (**D**) iSC neurons (squares or circles denote neurons also displaying build-up activity or not, respectively; same general format as Figure 2).

As previously mentioned, FEF inactivation had a stronger effect on visual neurons that also exhibited delay-or saccade-related activity. We found a similar pattern for the 113 ipsilesional iSC neurons with delay-period activity: 40 of 42 neurons exhibiting significantly reduced delay-period activity during FEF inactivation also exhibited either visual (62%), build-up (21%), or saccade-related activity (71%). Likewise, FEF inactivation consistently decreased visual (p < 0.001, z = 3.57, Wilcoxon signed rank test), and saccade-related activity prior to both visually-(p < 0.001, z = −3.59, Wilcoxon signed rank test) and memory-guided saccades (p < 0.05, z = −2.29, Wilcoxon signed rank test) in this subset of 40 neurons.

### FEF inactivation reduced saccade-related activity of ipsilesional iSC neurons

Next, we examined the FEF’s contribution to saccade-related activity in the ipsi-and contralesional iSC. For this analysis, it is imperative that saccades generated during FEF inactivation be matched as closely as possible for metrics and velocity, otherwise any changes in saccade-related activity could simply arise from the generation of a different saccade. We matched saccades within 1.5° of horizontal and vertical displacement and within 50°/s of radial peak velocity, although we typically found matches much less than these limits using a ranking procedure (see Materials and Methods; Figure 1F). Despite the generation of effectively equivalent saccades, FEF inactivation decreased ipsilesional saccade-related activity in ipsilesional iSC neurons. This result is shown for one neuron in Figure 4A, where activity for visually-guided saccades decreased significantly from 301 ± 85 to 263 ± 52 spikes/s (p < 0.001, z = 3.61, Wilcoxon signed rank test). Across our sample, FEF inactivation consistently decreased saccade-related activity for ipsilesional iSC neurons (minimum of 5 matches) for both visually-or memory-guided saccades (each p < 0.0001, z = −5.37 and −5.88, Wilcoxon signed rank tests, respectively, Figure 4B). The proportional decrease caused by FEF inactivation was greater for memory-versus visually-guided saccades, both in terms of how much activity decreased (20 ± 3% and 11 ± 2% decrease, respectively) and in the proportion of neurons exhibiting significantly changed activity (69% of 59 neurons, and 58% of 77 neurons, respectively).

**Figure 4.**
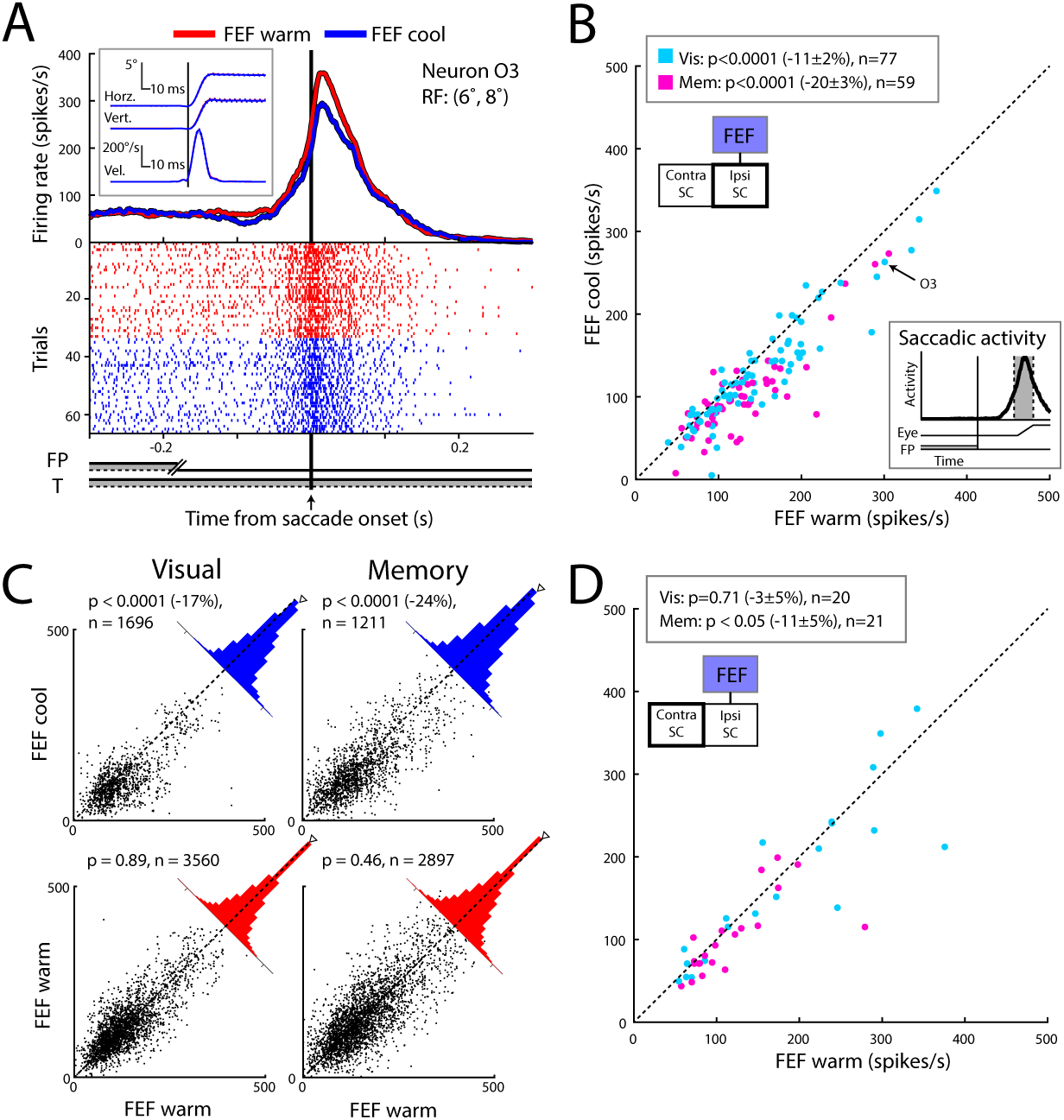
FEF inactivation decreased saccade-related activity of ipsilesional iSC neurons. (**A**) FEF inactivation decreased saccade-related activity in ipsilesional iSC neuron O3 (inset shows position and velocity profiles for matched visually-guided saccades). (**B**) FEF inactivation consistently decreased saccade-related activity (8 ms before saccade onset to 8 ms before saccade offset; see schematic) for ipsilesional iSC for both visually-and memory-guided saccades (same format as Figure 2B; neurons included only if they had at least 5 matched saccades). (**C**) Direct comparison of saccade-related activity for all matched contralesional saccades. FEF inactivation generally decreased saccade-related activity (top row; blue). As a control, we also matched saccades from FEF warm trials (bottom row; red), and did not find consistently decreased saccade-related activity. (**D**) FEF inactivation did not consistently influence saccade-related activity in the contralesional iSC.

Another way of analyzing the effects of FEF inactivation on saccade-related iSC activity is to directly compare activity for all matched saccades, pooled across all recorded neurons. The result of this analysis is shown in Figure 4C (top row), where each point represents saccade-related activity for a matched saccade generated during the FEF cool versus FEF warm condition. The clustering of points below the line of unity in the top row of Figure 4C, as well as the rightward skew of the blue histograms, reinforces how FEF inactivation decreases ipsilesional iSC saccade-related activity. As a control, we performed a similar saccade-matching procedure using FEF warm trials. While there is considerable scatter in this analysis around the line-of-unity (lower rows in Figure 4C), the resulting distributions of the red histograms were not skewed away from zero.

Finally, while FEF inactivation significantly decreased saccade-related activity in some contralesional iSC neurons (20 and 29% of neurons for visually-and memory-guided, respectively), such effects were less consistent across our sample (Figure 4D; 3± 5% decrease for visually-guided, p = 0.71, z = −0.37, 11 ± 5% decrease for memory-guided, p < 0.05, z = −2.14, Wilcoxon signed rank tests). Further, FEF inactivation decreased saccade-related activity significantly more for ipsi-versus contralesional iSC neurons during both the memory-guided saccade task (p < 0.05, z = −2.35, Wilcoxon rank sum test), but not the visually-guided saccade task (p = 0.07, z = −1.82, Wilcoxon rank sum test).

### Summary of effects of FEF inactivation on iSC activity

The results up to this point have focused on how FEF inactivation decreased visual, delay-period, and saccade-related activity in the downstream iSC, particularly in the ipsilesional side. In Figure 5, we summarize the impact of FEF inactivation on all aspects of functionally-defined activity in the ipsilesional iSC by presenting average spike density functions across our sample, separated where appropriate for visually-and memory-guided saccades. Presenting the data in this way emphasizes the greater impact of FEF inactivation on memory-guided saccades, which is consistent with previous behavioral (Peel et al., 2014) and neurophysiological (Sommer and Wurtz, 2000) findings of a larger contribution of FEF to working memory, and also eases comparisons with other studies (Koval et al., 2011; Johnston et al., 2014).

**Figure 5.**
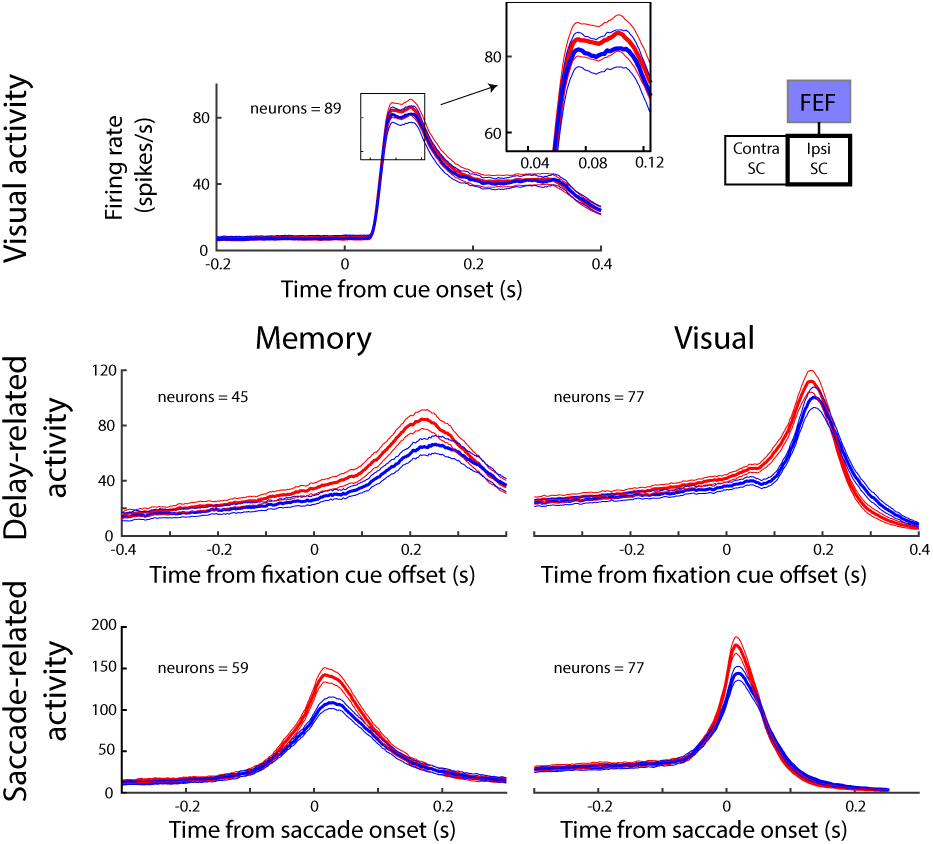
FEF inactivation reduced all aspects of activity in the ipsilesional iSC. For both visually-and memory-guided saccade tasks, FEF inactivation reduced the firing rate (mean ± SE) across ipsilesional iSC neurons possessing either visual, delay-period and/or saccade-related activity.

### Delays in the onset of iSC saccade-related accumulation predicted bilateral SRT increases

One of the intriguing features in Figures 4 and 5 is that saccade-related activity decreases during FEF inactivation, despite concomitant increases in SRT. Were SRT increases to be related to saccade threshold, then SC activity should have increased upon FEF inactivation. To reconcile these observations, we next examined how FEF inactivation altered the parameters of iSC pre-saccadic activity derived from a modified stochastic accumulator model (i.e., baseline and threshold activity, onset and rate of accumulation; see Materials and Methods), and determined which of these best predicted the accompanying SRT increases. Within this model, increased SRTs could arise from one or some combination of decreases in baseline activity or rate of accumulation, or increases in onset of accumulation or saccade threshold.

We first focus on the onset of accumulation. To find the onset of accumulation for each trial, we used a piecewise two-piece linear regression method (Figure 6A, see Materials and Methods for further details). This method systematically tests within a window for the inflection point between two linear regressions (one for baseline, and one for the rise in activity prior to saccade onset), with the onset of accumulation coinciding with the inflection point that minimizes the sum-of-square values for the two linear regressions against the convolved spike density function.

**Figure 6.**
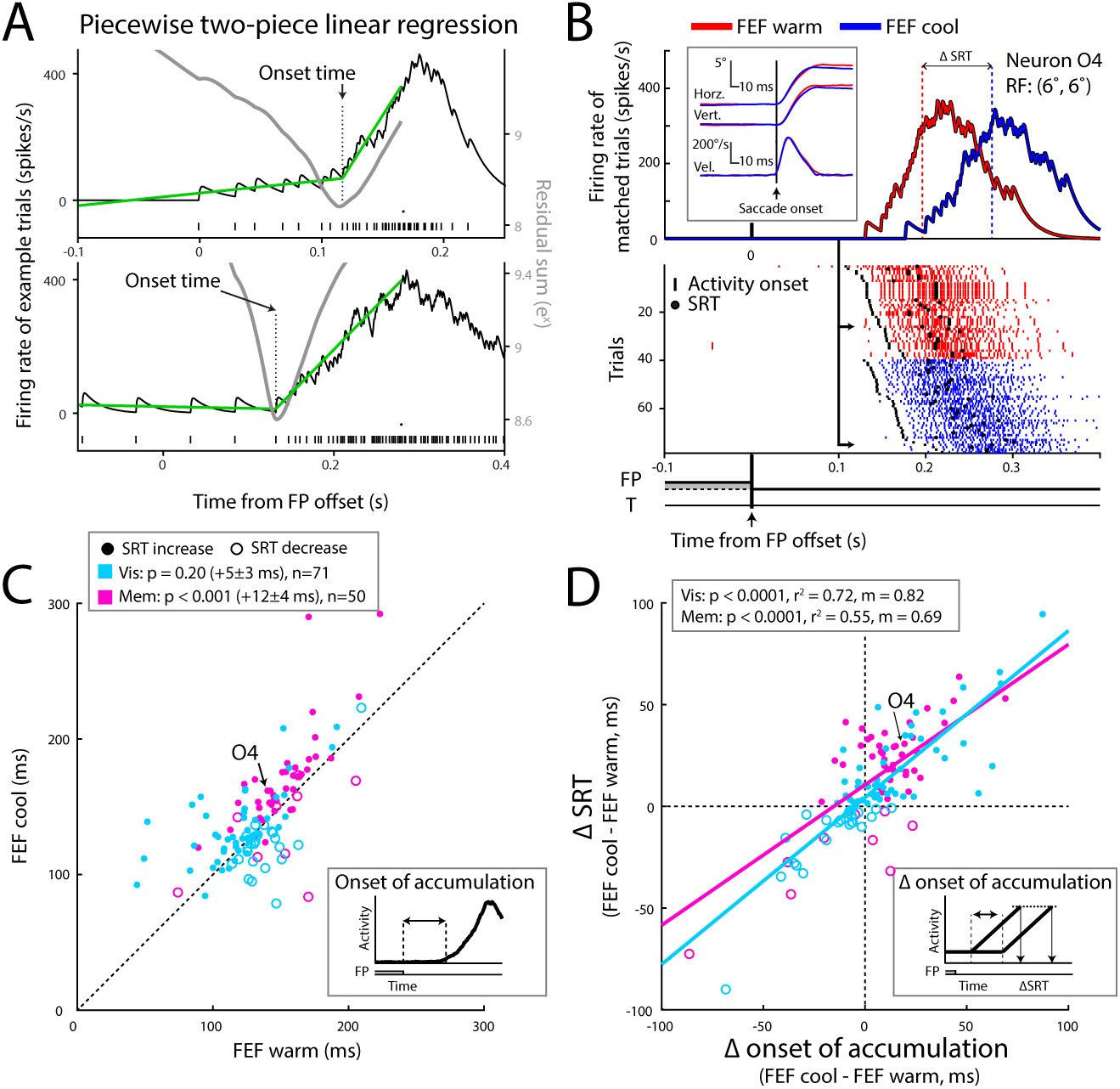
Inactivation induced changes in SRT correlated with delays in the onset of saccade-related activity in the ipsilesional iSC. (**A**) Depiction of how onset time is detected using a piecewise two-piece linear regression approach for two trials (see Materials and Methods for details). The onset time (dotted line) coincides with the inflection point that minimizes the summed squared error (grey curve, plotted against right axis) between convolved iSC activity (top, spike train shown below) and the two linear regressions (green lines). Note how the first linear regression captured any delay-period or buildup activity prior to the saccade (SRT is represented by the circle above raster plot). (**B**) For example ipsilesional neuron O4, FEF inactivation delayed the onset of saccade-related activity (ticks) and SRT (circles) for matched memory-guided saccades. Upper part shows spike density function and kinematics (inset) of one matched pair; lower part shows rasters for all matched saccades. (**C**) FEF inactivation generally delayed the onset of saccade-related activity in ipsilesional iSC neurons. Each point shows the average change per neuron (across trials with matched saccades), with filled or open circles denoting SRT increases or decreases, respectively. (**D**) Changes in the onset of saccade-related activity strongly correlated with concomitant changes in SRT.

The influence of FEF inactivation on the onset of accumulation (black ticks) and SRT (black circles) is shown for a representative neuron in Figure 6B, showing single-trial activity for one pair of matched saccades (top row Figure 6B; inset shows the position and velocity profiles for this match), and across all matches recorded from this neuron (Figure 6B; bottom). Across all matched trials for this neuron, FEF inactivation increased the average onset of accumulation from 142 ± 16 to 162 ± 22 ms after fixation cue offset (p < 0.0001, z = −3.91, Wilcoxon signed rank test), and also significantly increased the average SRT of matched memory-guided saccades from 207 ± 20 to 238 ± 29 ms (p < 0.0001, z = −4.39, Wilcoxon signed rank test). Thus, in this example, FEF inactivation delayed onset of iSC accumulation and increased SRT.

Across our sample, FEF inactivation delayed the onset of iSC accumulation in the ipsilesional iSC for memory-guided (Figure 6C; 12 ± 3 ms increase, p < 0.001, z = 3.53, Wilcoxon signed rank test; onsets from individual trials were averaged within neurons for this analysis) saccades, but not visually-guided saccades (5 ± 3 ms increase, p = 0.20, z = 1.29, Wilcoxon signed rank test). We surmise that the lack of a significant increase for visually-guided saccades may relate both to the smaller accompanying changes in SRT with this task, and to our stringent saccade-matching analyses, as significant increases in accumulation onset were observed in the absence of saccade matching for both visually-guided (increases of 16 ± 3 ms, p < 0.0001, z = 4.82, Wilcoxon signed rank test), and memory-guided saccades (increases of 17 ± 3 ms, p < 0.0001, z = 4.81, Wilcoxon signed rank test). More importantly, as shown in Figure 6C, increases or decreases in the onset of accumulation during FEF inactivation generally corresponded to a similar increase (e.g., closed circles clustering above the line of unity in Figure 6C) or decrease (e.g., open circles clustering below the line of unity in Figure 6C) in SRT, respectively. To analyze this more closely, we plotted the change in the onset of accumulation versus the change in SRTs on a neuron-by-neuron basis, and determined the variance explained by a linear correlation (Figure 6D). Interestingly, we found robust correlations across ipsilesional iSC neurons for both visually-and memory-guided saccades (r^2^ values of 0.72 and 0.55, p values < 0.0001 for each, F statistics of 175.28 and 58.64, respectively), with slopes near 1.0 (0.82 and 0.69, respectively) and intercepts near 0 ms (4 and 11 ms, respectively). Notably, we also observed similar patterns in how changes in the onset of accumulation within the contralesional iSC predicted the accompanying changes in SRT (r^2^ = 0.44, p < 0.01, F = 12.83 for visually-guided saccades, and r^2^ = 0.60, p < 0.001, F = 23.68 for memory-guided saccades). We verified our findings using various approaches to detect onset time (see Methods and Materials), including an alternative Poisson-based analysis. Results using this Poisson-based analysis agreed with the two-piece linear regression method, with FEF inactivation delaying the onset of accumulation in the ipsilesional iSC (increase of 6 ± 4 ms, p = 0.13, z = 1.51, n = 71 for visually-guided, and increase of 11 ± 5 ms, p < 0.0001, z = 4.15, n = 54 for memory-guided saccades, Wilcoxon signed rank tests), and the changes in onset remained strongly correlated with SRT differences (r^2^ = 0.53, p < 0.0001, F = 78.41, m = 0.53, y-int = 6 for visually-guided saccades, and r^2^ = 0.64, p < 0.0001, F = 93.56, m = 0.61 y-int = 11 for memory-guided saccades). Thus, on a neuron-by-neuron basis, changes in the onset of accumulation of saccade-related iSC activity predicted accompanying changes in SRT in an almost one-to-one manner.

### FEF inactivation altered other parameters in the ipsilesional iSC neurons, but such changes did not predict accompanying SRT increases

SRT increases with FEF inactivation could also be related to increases in threshold activity, or decreases in baseline activity or rate of accumulation. While FEF inactivation did impact these parameters, the variability or direction of changes in neural activity did not relate as well to accompanying changes in SRT. For instance, FEF inactivation decreased the accumulation rate (4.6 ± 1.1 to 3.5 ± 1.1 spikes/s^2^, p < 0.0001, z = 4.70, Wilcoxon signed rank test), and threshold activity (210 ± 40 to 179 ± 48 spikes/s, p < 0.01, z = 2.90, Wilcoxon signed rank test) in an exemplar neuron (see arrow in Figure 7A; same neuron as in Figure 6B). Across our sample of ipsilesional iSC neurons, FEF inactivation consistently decreased baseline (Figure 7B, decrease of 2 ± 1 spikes/s, p < 0.01, z = −2.60, Wilcoxon signed rank test) and threshold activity (Figure 7C, decrease of 11 ± 3%, p < 0.001, z = −3.34, Wilcoxon signed rank test) prior to memory-guided saccades, but not visually-guided saccades (p values = 0.22 and 0.13, z = −0.29 and −1.52, Wilcoxon signed rank tests for baseline and threshold activity, respectively). FEF inactivation did not influence the rate of accumulation in either task (Figure 7D; p values = 0.76 and 0.06, z = −0.30 and – 1.85, Wilcoxon signed rank tests for visually-and memory-guided tasks, respectively). Although changes in the rate of accumulation or baseline would be consistent with a neuronal mechanism that increases SRT during FEF inactivation, it is also important to consider interactions among parameters, and how these collective changes could quantitatively relate to SRT increases.

**Figure 7.**
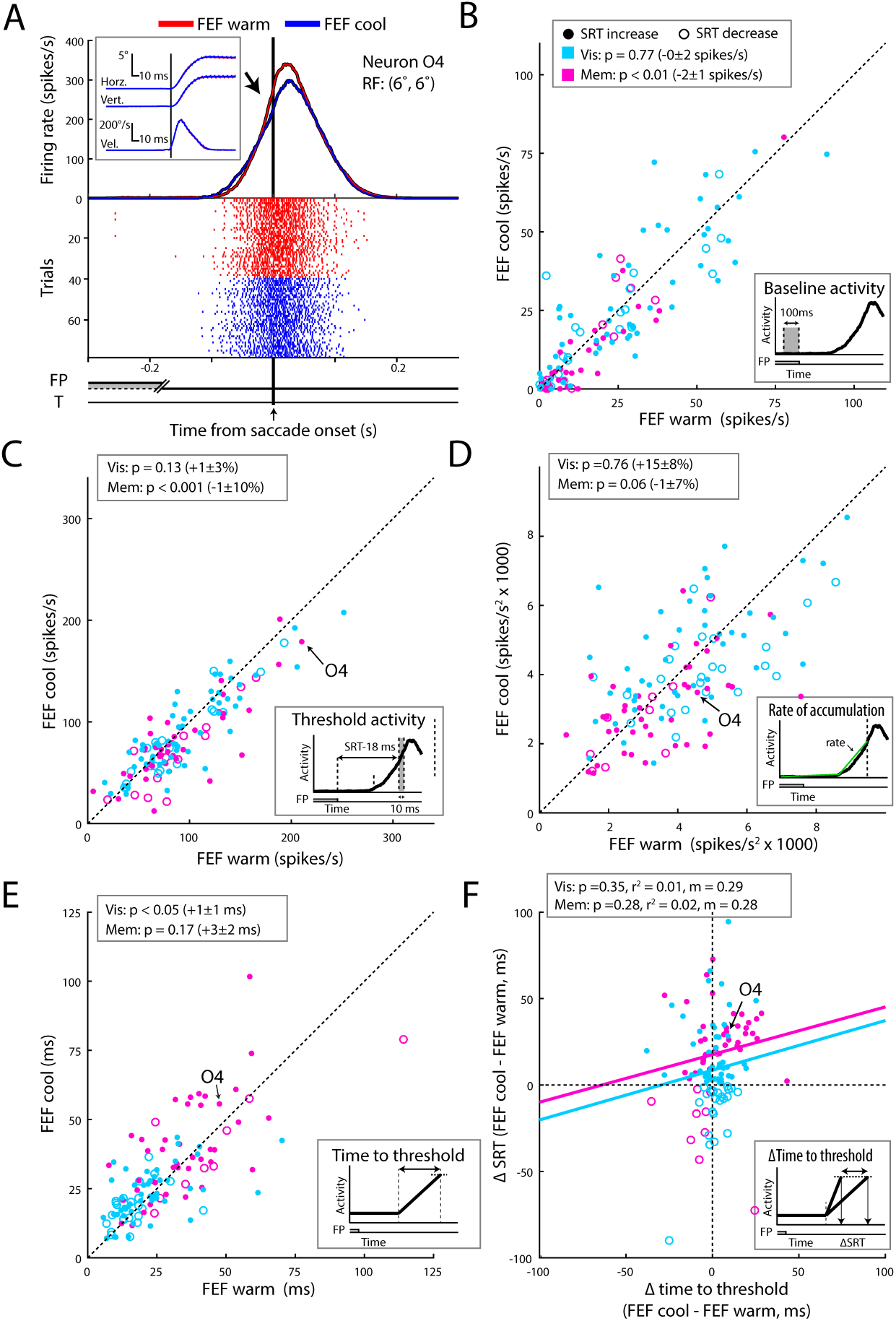
FEF inactivation changed other aspects of iSC activity, but these changes poorly predicted changes in SRT. (**A**) In example neuron O4, FEF inactivation decreased both the accumulation rate and threshold activities (see Materials and Methods for how these parameters were measured) for matched memory-guided saccades. Same format as Figure 6B. **(B-D)** Across our sample, FEF inactivation decreased baseline activity (**B**), threshold activity (**C**), accumulation rate (**D**) of ipsilesional iSC neurons, particularly for memory-guided saccades. (**E, F**) To analyze how changes in these parameters related to differences in a one-to-one manner, we computed the time to reach threshold as the difference of threshold and baseline activities divided by the accumulation rate. Note how this parameter does not directly incorporate the onset of accumulation, so it assumes that activity starts to accumulate at an arbitrary point in time after the go-cue. While such changes did increase the time to reach threshold in the ipsilesional iSC (**E**), such changes did not fully account for the concomitant changes in SRT (**F**).

To examine how well such parametric changes related to accompanying changes in SRT, we employed a standard accumulator model that allowed us to extract the *time* to *reach threshold*, computed from the measured values for baseline, threshold, and accumulation rate ([threshold-baseline activity]/[accumulation rate]; see Materials and Methods). Accumulator models are most commonly applied to speeded, rather than delayed, response tasks, but nevertheless such models provide a framework in which to relate changes in iSC activity to accompanying changes in SRT. Note that this time to reach threshold value does not directly incorporate the onset of accumulation; effectively this model assumes that neural activity accumulates from baseline at some fixed time after a go cue. The results of this analysis are shown on a neuron-by-neuron basis in Figure 7E and F. While the time to reach threshold in the ipsilesional iSC marginally increased with FEF inactivation for visually-guided saccades (1 ± 1 ms increase, p < 0.05, z = 2.25, Wilcoxon signed rank test), we observed no consistent effects on memory-guided saccades (p = 0.17, z = 1.37, Wilcoxon signed rank test), and a linear regression of such changes against concomitant changes in SRT revealed no correlation in the ipsilesional iSC for either visually-(p = 0.35, F = 0.89, r^2^ value of 0.01) or memory-guided saccades (p = 0.28, F = 1.20, r^2^ value of 0.02). Moreover, any changes in the time to reach threshold for the contralesional iSC did not relate at all to associated changes in ipsiversive SRT (p = 0.55 and 0.97, F = 0.38 and 0.001 for visually-and memory-guided saccades, respectively). Thus, even when incorporating the three parameters of baseline, threshold, and accumulation rate, such changes do not fully account for the accompanying changes in SRT in the ipsilesional iSC.

### Changes in the onset of accumulation best explained SRT differences across matched saccades, even without FEF inactivation

We extended the above analyses to the level of individually-matched saccades, which enabled us to test how changes in onset time correlated with a larger distribution of SRT changes. For each matched saccade pair, extracted either across FEF inactivation (top-left subplot in Figure 8A) or only from trials without FEF inactivation (bottom-left subplot in Figure 8A) from the ipsilesional iSC, we derived both a change in the onset of accumulation and a change in SRT. We pooled trials across saccade tasks, since we found largely equivalent results for visually-or memory-guided saccades (r^2^ values of 0.62 and 0.42, F = 1731.2 and 622.8, respectively, for relationship of changes in onset time versus SRT differences with FEF inactivation). Note that this procedure often matches trials where SRT decreased upon FEF inactivation, due to the overlap between RT distributions when the FEF was or was not inactivated. Regardless, the changes in SRT and onset of accumulation remained highly correlated, regardless of whether the matched pairs were being compared across FEF inactivation or not (p values less than 0.0001, F = 2201.4 and 4684.1, r^2^ values of 0.53 and 0.51, slopes of 0.65 and 0.59, and y-intercepts of 8 and 0 ms, respectively). In contrast, comparison of the change in the time to reach threshold (computed from the baseline, rate of accumulation, and threshold) versus the change in SRT for matched-saccade pairs revealed much weaker relationships (right columns in Figure 8A). Moreover, we found similarly poor relationships from a time to reach threshold computed from a completely independent measure of accumulation rate (r^2^ values of 0.10 and 0.11 for with or without FEF inactivation, respectively, see Materials and Methods).

**Figure 8.**
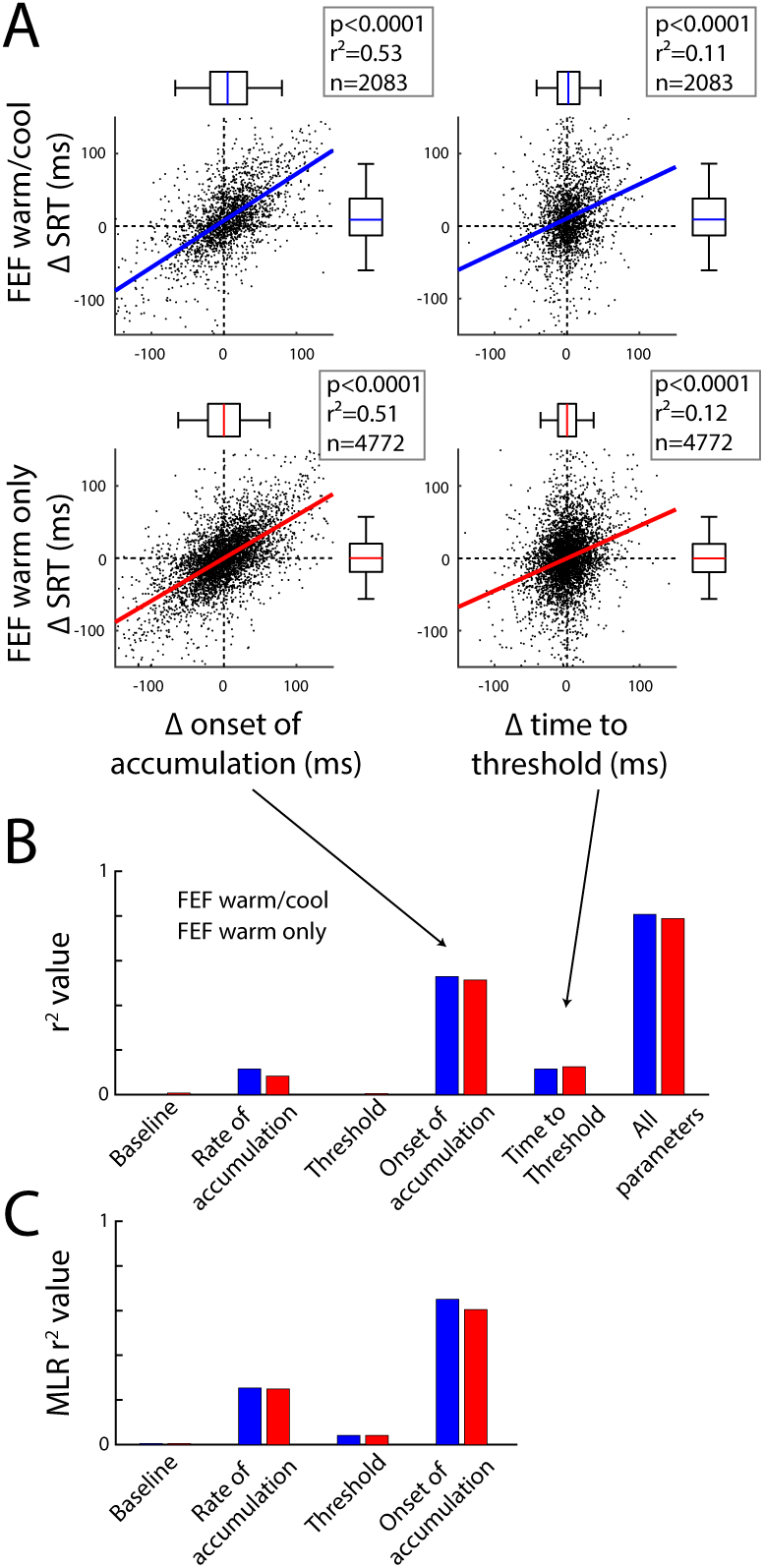
Across all matched trials, changes in the onset of accumulation best reflected changes in SRT, even without FEF inactivation. (**A**) Across all matched trials extracted with (top row, blue lines) and without FEF inactivation (bottom row, red lines), differences in the onset of accumulation (left column) in the ipsilesional iSC related better to associated changes in SRT, as compared to differences in the time to reach threshold (right column). (**B**) Amount of SRT variance explained by different combinations of individual or grouped parameters extracted from a rise-to-threshold model. Regardless of whether matched pairs were extracted across FEF inactivation or not, consideration of the change in the onset of accumulation greatly increased how well changes in iSC activity predicted concomitant changes in SRT. (**C**) Across trials matched with or without FEF inactivation, the onset of accumulation best correlated with the remaining residual error following a multiple linear regression of the other individual parameters and SRT.

We repeated these analyses independently for each of the parameters of baseline, rate of accumulation, or threshold, and as shown by r^2^ values in Figure 8B, changes in these single parameters were also very poor predictors of accompanying changes in SRT. In contrast, consideration of the onset of accumulation either alone or in conjunction with all other parameters (i.e., adding the onset time of accumulation to the time to reach threshold) greatly increased the relationship between changes in iSC activity and changes in SRT (Figure 8B). Perhaps just as importantly, these observations held regardless of whether the matched pairs were compared across FEF inactivation or not (blue or red bars in Figure 8B, respectively), or from contralesional iSC (**data not shown**).

Finally, to address how well a single parameter could independently explain SRT changes, even when allowing for correlation among other parameters, we adopted a multiple linear regression approach (Hewitt et al., 2015) within the framework of a rise-to-threshold model of saccade initiation (see Materials and Methods). The objective of this analysis was to determine how much an individual parameter could independently explain the remaining variance following a multiple linear regression that correlates all the other parameters to SRT. As shown in Figure 8C, the onset of accumulation best accounted for the remaining variance from a multiple linear regression consisting of the other three parameters, reaffirming the importance of this parameter.

### Summary of Results

Reversible inactivation of a large volume of the unilateral FEF decreased all aspects of ipsilesional iSC activity, consistent with a general loss of excitatory input. The magnitude of such decreases in iSC activity depended both on the functional content of the signal, with greatest decreases for saccade-related activity, and on the inferred depth of the neuron within the SC, with greater decreases in visually-related activity on those neurons also displaying delay-and saccade-related activity. Such results largely conform both with the preferential distribution of frontal projections to intermediate and deeper layers of the iSC (Tigges and Tigges, 1981), and with antidromic studies of the functional content of corticotectal neurons (Segraves and Goldberg, 1987; Sommer and Wurtz, 2000; Helminski and Segraves, 2003). The absence of any effect of FEF inactivation on the latency of the first spike following visual stimulus onset is also consistent with evidence that visual responses in the iSC require the integrity of the retinogeniculostriate pathway passing through the magnocellular laminae of the LGN (Schiller et al., 1979); perhaps not too surprisingly given the short response latencies, the FEF is also not a critical node in the pathway mediating the initial timing of the visual response in the iSC. Finally, by fitting the profiles of saccade-related iSC activity to a stochastic accumulator model, we showed that delays in the onset of saccade-related accumulation in either iSC, rather than systematic changes in either saccade threshold, baseline, or the rate of accumulation, best-explained concomitant changes in SRT in the delayed saccade task, emphasizing the need to consider the onset of accumulation as a parameter in neurophysiologically-inspired models of saccade initiation.

## Discussion

### FEF inactivation reduces excitatory input to the ipsilesional iSC without disinhibiting the contralesional iSC

The functional content of cortical signals relayed directly to iSC has been well-characterized using antidromic identification (Segraves and Goldberg, 1987; Everling and Munoz, 2000; Sommer and Wurtz, 2000; Wurtz et al., 2001; Helminski and Segraves, 2003). However, due to other polysynaptic pathways such as through the basal ganglia or other cortical areas, inactivation studies like ours are required to causally assess the collective influence of the FEF on iSC activity. The general agreement between our results and those that would have been predicted by antidromic studies alone is encouraging, reaffirming the FEF’s role in providing excitatory input to the ipsilateral iSC, particularly for those neurons displaying saccade-related activity.

Long-range interactions between different regions of the iSC or between the FEFs are thought to be inhibitory (Munoz and Istvan, 1998; Schlag et al., 1998). Could the general reduction in ipsilesional iSC activity during FEF inactivation, and the associated increases in RT, arise from disinhibition of the contralesional FEF and/or iSC, or a shift toward increased activity of fixation neurons in the rostral iSC (Munoz and Wurtz, 1993; Dorris and Munoz, 1995)? While we did not record from the rostral iSC or contralesional FEF, a number of observations argue against these interpretations. First, the magnitude of contralesional iSC activity neither increased nor decreased, contrary to what would have been expected from disinhibition or increased rostral iSC activity, respectively. Second unilateral FEF inactivation decreased the peak velocity and prevalence of cue-related microsaccades in both directions (Peel et al., 2016), which is more consistent with decreasing, rather than increasing, levels of rostral iSC activity (Hafed et al., 2009). Third, although FEF inactivation decreased the magnitude of visual responses on ipsilesional saccade-related iSC neurons, the latency of such responses was unchanged (Figure 2). In contrast, during paradigms associated with increased rostral iSC activity, visual responses in the caudal iSC are both reduced in magnitude and delayed in onset latency (Marino et al., 2012).

Although FEF inactivation reduced ipsilesional iSC activity and delayed the onset of accumulation across our sample, there were instances where FEF inactivation had little to no effect on ipsilesional iSC activity, or decreased rather than increased the onset of accumulation. There are a number of potential reasons for this variability. It may be that not every iSC neuron we recorded receives input from the FEF, direct or otherwise. Further, any impact of FEF inactivation may be masked by compensatory inputs from other structures that converge on the same neuron. To our knowledge, there is no data that speak to this level of anatomical and physiological detail within the oculomotor network. Additionally, given the topographically-aligned nature of projections from the FEF to the iSC (Sommer and Wurtz, 2000), it is possible that we occasionally recorded from iSC locations that were not impacted by FEF inactivation. The question of spatial register may also explain why FEF inactivation occasionally decreased both SRTs and the onset of accumulation (e.g., the points in the lower-left quadrant in Figure 6D). We did not find any spatial dependency in how much FEF inactivation impacted iSC activity or behavior (e.g., larger cooling effects did not stem from particular parts of the visual field), but this analysis was hampered by the variability in loop location across monkeys and hemispheres.

### A bilateral influence of the unilateral FEF on SRT and the onset of saccade-related accumulation in the iSC

The failure to observe disinhibition in the contralesional iSC is surprising given that focal pharmacological inactivation of the FEF facilitates ipsiversive oculomotor behaviors (Sommer and Tehovnik, 1997; Dias and Segraves, 1999; Wardak et al., 2006). Studies using focal microstimulation (Schlag et al., 1998; Seidemann et al., 2002) or paired bilateral recordings (Cohen et al., 2010) have also supported a view wherein the two FEFs compete in a push-pull fashion. In contrast, large-volume FEF inactivation tends to delay rather than facilitate ipsiversive oculomotor behaviors (Figure 1; (Peel et al., 2014, 2016; Kunimatsu et al., 2015)), and although ipsiversive SRT increases were of lower magnitude and more idiosyncratic than the increases in contraversive SRTs, when present such SRT increases related best to delays in the onset of saccade-related accumulation in the corresponding iSC.

The behavioral and neurophysiological results produced by unilateral cryogenic inactivation of the FEF, reported here and elsewhere (Peel et al., 2014, 2016), also differ from that seen following unilateral cryogenic inactivation of the adjacent dorso-lateral prefrontal cortex (DLPFC). As reported by (Koval et al., 2011; Johnston et al., 2014), unilateral cryogenic inactivation of the DLPFC via cooling loops implanted within the caudal principal sulcus shortens the RTs of ipsiversive saccades and increases preparatory, visual-, and saccade-related activity in the contralesional iSC. Thus, unlike what we observed following unilateral inactivation of the FEF, unilateral cryogenic inactivation of the nearby DLPFC does produce results consistent with disinhibition via a push-pull mechanism. Importantly, since the cooling loops implanted in the FEF or DLPFC were placed within the arcuate or caudal principal sulci respectively, it is unlikely that they inactivated overlapping volumes of tissue on the gyral crowns.

### New perspectives on saccade initiation

FEF inactivation altered how iSC activity relates to saccade initiation, doing so in a manner that provides new insights into how the oculomotor brainstem initiates a saccade. One fundamental observation is that less saccade-related activity is emitted by ipsilateral iSC neurons during FEF inactivation, even for saccades matched closely for metrics and kinematics. Clearly, at the level of single iSC neurons, saccade threshold can vary for movements to the center of the response field (Figure 7). This observation complements observations of changes in saccade threshold in the iSC (Everling et al., 1999; Jantz et al., 2013) or FEF (Heitz and Schall, 2012) during different tasks or cognitive sets. Heitz and Schall (2012) reconciled observations of increased SRTs despite decreasing thresholds by proposing a leaky integrator mechanism where saccade-related spikes are integrated over time to produce an invariant level of cumulative activity, and our results of decreasing thresholds and rates of accumulation upon FEF inactivation show that a similar mechanism may apply to the iSC. The decreased activation of iSC neurons for movements to the center of the response field could also be offset by increased activity from other off-center iSC neurons, extending the notion of saccade threshold within the iSC to forms of population coding envisaged for other aspects of saccade control (see Gandhi and Katnani, 2011 for review). It is also possible that an overall decrease in iSC activity could be offset by increasing activity in other areas that project to the brainstem burst generator, such as the fastigial nucleus (Noda et al., 1990), to produce an overall equivalent input to the brainstem burst generator. Understanding the contribution of these other cell populations, either within or outside of the iSC, is presumably required to fully explain SRT changes with FEF inactivation.

Previous evidence for fixed thresholds in the FEF (Hanes and Schall, 1996; Brown et al., 2008) and iSC (Paré and Hanes, 2003) fit well with rise-to-threshold (Carpenter and Williams, 1995; Reddi and Carpenter, 2000; Lo and Wang, 2006; Carpenter et al., 2009) and drift-diffusion models (Ratcliff et al., 2003, 2007) of saccade initiation, strengthening contentions that these models provided useful descriptions of neural activity. However, by embracing a greater subset of experimental tasks or conditions, the work of Heitz and Schall (2012, 2013) and Jantz and colleagues (2013) revealed that contemporary stochastic accumulator models failed to predict observed profiles of FEF or iSC activity. More broadly, delays in the onset of movement-related activity in the FEF and/or iSC, rather than changes in threshold or rate of accumulation, relate best to SRT increases in more difficult visual search paradigms (Woodman et al., 2008), or to post-error increases in SRT within a stop-signal paradigm (Pouget et al., 2011). A recent review has emphasized the importance of incorporating the onset of activity within studies of decision-making, particularly in tasks requiring top-down regulation (Teichert et al., 2016). Our inactivation findings, as well as our comparison of matched saccades even without FEF inactivation, extend these findings by demonstrating the importance of the onset of saccade-related activity in the iSC in delayed response tasks, and by showing that such onset is governed at least in part by inputs from the FEF. Taken together, it is becoming increasingly clear that the onset of saccade-related activity is a relevant metric that impacts SRT, although future studies incorporating FEF inactivation within speeded response tasks are required to see whether our results generalize more widely.

## Acknowledgements

This work was supported by operating grants from the Canadian Institutes of Health Research to BDC (MOPs: 93796, 123247 and 142317) and the Natural Sciences and Engineering Research Council (NSERC; RGPIN-311680). TRP was supported by an Ontario Graduate Scholarship and TRP and SD were supported by funding from an NSERC CREATE grant.

